# Epithelial regulatory networks link non-coding genetic variation to clinical heterogeneity in ulcerative colitis

**DOI:** 10.64898/2026.08.04.742507

**Authors:** Yufan Liu, John P Thomas, Balazs Bohar, Nick Powell, Dezso Modos, Alexandra Paun, Tamas Korcsmaros

## Abstract

Ulcerative colitis (UC) is a genetically heterogeneous disease causing chronic intestinal inflammation. Genome-wide association studies have linked numerous non-coding single nucleotide polymorphisms (SNPs) to UC, yet how these variants relate to intestinal epithelial function and clinical outcomes remains largely unknown. To address this, we aimed to reconstruct patient-specific epithelial signalling and regulatory networks perturbed by non-coding SNPs and to dissect patient heterogeneity in a large UC phase III trial. We analysed genotype data from 452 participants in the etrolizumab phase III HICKORY cohort using the integrated SNP network platform (iSNP) to map the downstream effects of non-coding SNPs and identify transcription factors (TFs) they perturb in each patient. TFs were filtered to retain those with differential regulatory activity in UC epithelial cells, patients were stratified by these TF profiles, and clusters were correlated with etrolizumab outcomes. We identified 14 SNP-propagated TFs with differential epithelial regulatory activity. Unsupervised clustering stratified patients into subgroups differing in disease severity and treatment response: two clusters had the highest severity and lowest remission and response rates, whereas two others had the best outcomes. iSNP-based clustering improved endoscopic healing prediction (*p* = 0.03), with trends for clinical response and histological endoscopic mucosal improvement. This systems genomics framework links patient-specific non-coding SNPs to epithelial signalling and regulatory networks in UC and captures clinically meaningful heterogeneity associated with treatment outcomes. It supports genetics-informed precision medicine and improved clinical trial design in inflammatory bowel disease and beyond.

## Introduction

Ulcerative colitis (UC), a major clinical subtype of inflammatory bowel disease (IBD), arises due to complex interactions between genetic, immune, environmental, and microbial factors [1]. The multifaceted nature of UC pathogenesis, together with substantial patient heterogeneity, poses significant challenges for clinical management. Whilst there has been considerable progress in expanding the therapeutic options available for patients in recent years, a major therapeutic ceiling remains with only 30% of patients achieving durable clinical remission [2]. Even in rigorous randomised controlled trials (RCT), treatment effects over placebo are modest, ranging from ∼30% to as low as ∼7% for currently licensed therapies [3].

Data from clinical trials and real-world studies have consistently demonstrated patient heterogeneity in treatment responses, but the mechanisms underpinning this variability remain poorly characterised. The era of genome-wide association studies (GWAS), which delivered remarkable success in identifying more than 240 genetic loci associated with IBD [4], has yet to deliver on the promise of genomics-informed precision medicine in IBD. A major challenge is the predominance of IBD-associated single nucleotide polymorphisms (SNPs) within non-coding genomic regions that are challenging to functionally annotate [5]. Moreover, GWAS have largely failed to identify reproducible susceptibility loci associated with therapeutic response to advanced therapies in IBD, with the exception of variants underpinning immunogenicity to anti-TNF agents [6]. Whilst other omic layers have offered attractive candidates, such as oncostatin M (OSM) through transcriptomics analysis [7], or short-chain fatty acids and bile acids highlighted in metabolomics studies [8], none have yet yielded clinically implemented predictors of treatment response.

This lack of biomarkers for stratifying patients has hindered the successful translation of many promising drug candidates identified in pre-clinical models into effective clinical treatments when evaluated in human clinical trials. Mechanistic pre-clinical studies, which often rely on single genetic backgrounds such as murine models, fail to capture the heterogeneity of human disease. As a result, they may pinpoint druggable pathways that are active only in specific subsets of patients. When tested in heterogeneous patient populations, such therapies may fail to achieve significant differences in efficacy compared to placebo across the entire patient cohort, despite genuine efficacy in certain subgroups.

One recent example is the unexpected failure of etrolizumab to demonstrate consistent efficacy across multiple phase III RCTs in IBD, despite promising pre-clinical and early-phase clinical trial data [9]. Etrolizumab is a dual-action anti-β7 monoclonal antibody that selectively targets α4β7 and αEβ7 integrins, which modulates both the trafficking of immune cells into the gut and their inflammatory effects on the intestinal epithelium. In the HICKORY phase III RCT, etrolizumab achieved statistical significance for induction of clinical remission compared to placebo (18.5% vs 6.3%, p=0.0033) but failed to meet the primary maintenance endpoint (24.1% vs 20.2%, p=0.50) in moderate-to-severe UC patients previously treated with anti-TNF therapy [10]. Similar results were observed in the LAUREL trial, which evaluated the efficacy of achieving maintenance in UC patients without prior anti-TNF failure [11], while the HIBISCUS I and II induction trials also yielded mixed results [12]. These failures and inconsistent trial outcomes underscore the urgent need for new patient stratification methods that can resolve pathogenic and treatment heterogeneity in IBD.

Genomics-based stratification represents a particularly attractive strategy to address this gap. Germline variants are stable across the life course and reproducible across independent cohorts, unlike other omic layers such as transcriptomic, metabolomic, or metagenomic readouts, which are highly context dependent and influenced by environmental exposures. To address this, we developed the integrated single nucleotide polymorphism network platform (iSNP), a systems genomics framework that maps non-coding SNPs to regulatory genomic regions, including transcription factor (TF) binding sites and microRNA target sites, and propagates these perturbations through signalling and gene-regulatory networks (GRNs) [13]. Unlike conventional GWAS, iSNP integrates non-coding SNP effects into patient-specific signalling and regulatory networks, and through unsupervised clustering of patients reveals subgroups defined by distinct pathogenic mechanisms. Notably, our recent analysis highlighted the central role of transcriptional regulation in capturing patient heterogeneity and potential treatment responses in IBD [14].

Here, we harnessed iSNP to dissect patient heterogeneity in the HICKORY cohort by identifying genotype-driven gene regulatory mechanisms that may explain variability in treatment outcomes to etrolizumab. Given etrolizumab inhibits the recruitment of T cells to the intestinal mucosa and the retention of intra-epithelial lymphocytes in the epithelial compartment [15], we contextualised this analysis to the colonic epithelium. Moreover, the intestinal epithelium is a prime compartment where genetic risk may manifest in IBD [16], acting as the physical and immunological interface with the microbiota, integrating cytokine and environmental cues, and its integrity correlates with outcomes such as mucosal healing [17]. Furthermore, epithelial barrier dysfunction has been observed in both active and quiescent IBD and is linked to relapse risk and adverse outcomes [18]. However, how non-coding SNPs disrupt epithelial signalling networks and downstream gene-regulatory programmes, and how heterogeneity in these disrupted networks relates to IBD severity and treatment outcomes remain poorly defined.

To address this gap, we modelled the cumulative impact of patient-specific non-coding SNPs on epithelial cell–specific signalling processes in the HICKORY cohort. We then integrated these networks with epithelial gene-expression data to infer TF activity and to characterise patient subgroups exhibiting distinct patterns of epithelial regulatory rewiring. By linking these TFs to patient-specific clinical outcomes in the HICKORY cohort, we provide mechanistic insights into how non-coding SNPs shape pathogenic heterogeneity and influence therapeutic response. To our knowledge, this represents the first study connecting patient-specific genotype-driven regulatory network heterogeneity to robustly characterised clinical trial outcomes in UC.

## Results

### Differential gene expression reveals genes associated with treatment remission and failure to etrolizumab

To characterise the molecular features associated with week 14 clinical outcomes to etrolizumab, we first performed differential expression analysis comparing clinical remission with non-remission using colonic whole genome bulk transcriptomics data from patients at both baseline (week 0, n = 447) and post-induction (week 14, n = 399). Across these contrasts, we used a robust threshold of absolute log2 fold-change > 1.0 and adjusted p < 0.01 to indicate differential gene expression. The transcriptional landscape associated with remission and non-remission to etrolizumab in baseline biopsies is summarised in Figure 1a.

**Figure 1.**
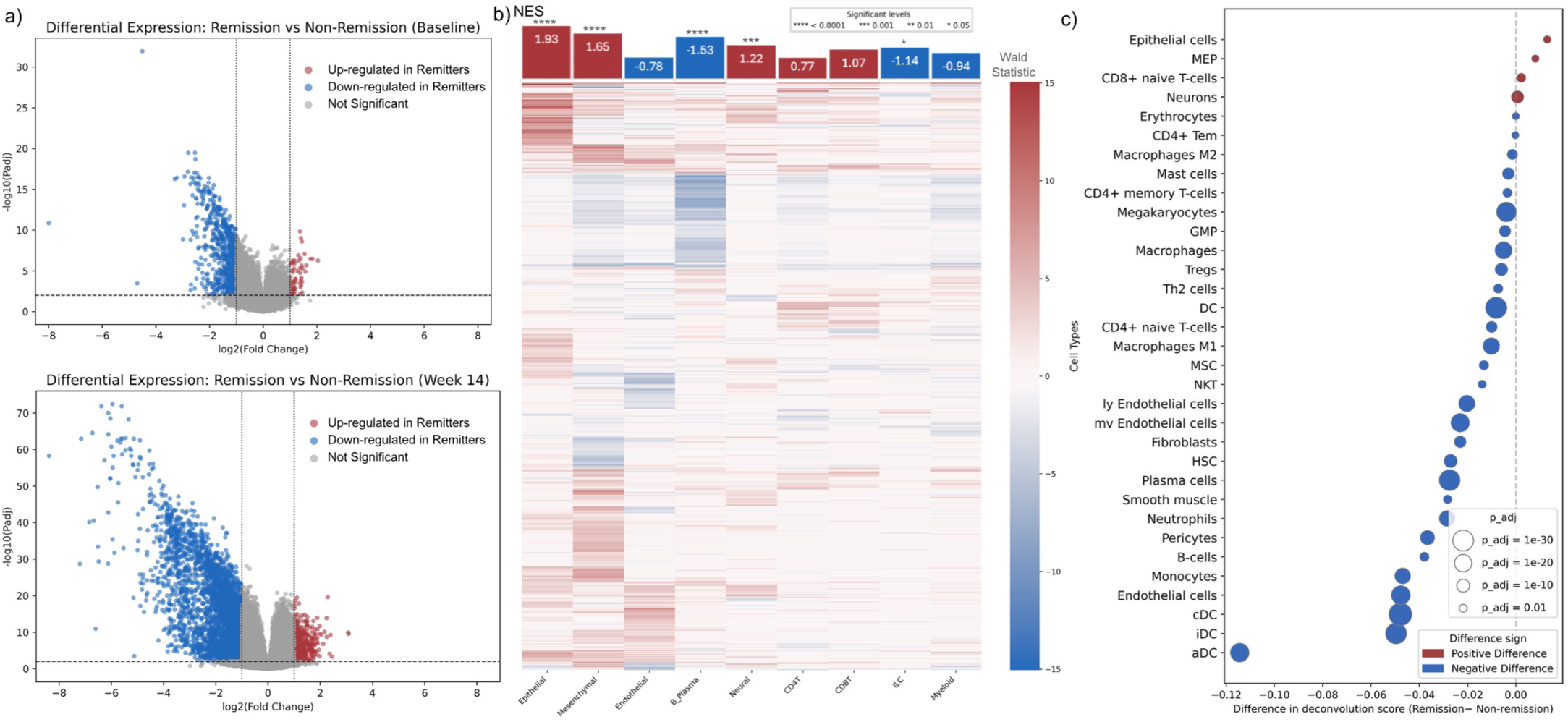
Differential Gene Expression and Cellular Composition in Ulcerative Colitis Remission. a) Volcano plot of differential expressed genes between clinical remission and non-remission at baseline (upper) and at week 14 (lower); b) This heatmap shows genes (y-axis) that are differentially expressed between remission and non-remission across cell types (x-axis). Each tile reports the DESeq2 statistics for UC-inflamed versus healthy controls for that gene within that cell type. Red indicates positive values (higher in UC-inflamed), blue indicates negative values (lower in UC-inflamed); The bar plot on the top shows the normalised enrichment score (NES) indicating the enrichment level of remission-associated genes in each cell type. Red indicates positive NES and blue indicates negative NES, starts represents the statistically significant level; c) Cell types with statistically significant differences in xCell cell deconvolution scores between remission and non-remission patients at week 14, restricted to padj < 0.05. The colour of each point denotes the direction of the difference (red for remssion and blue for non-remission s). Point size is proportional to-log10(padj).

To further delineate the cellular and biological processes underlying these transcriptional differences, we mapped the differentially expressed genes (DEGs) to reference cell-type signatures from the single cell IBD (scIBD) meta-analysis resource [19]. This analysis revealed a marked enrichment of remission-associated DEGs within epithelial cell–associated gene programmes, suggesting that the epithelial compartment contributes substantially to the molecular differences between clinical remission and non-remission in etrolizumab-treated UC patients (Figure 1b).

We next performed cellular deconvolution analysis of the bulk transcriptomics data using xCell [20]. Comparison of xCell cell type enrichment scores between patients in remission and non-remission at week 14 revealed that only epithelial and MEP (microfold/enterocyte progenitor) populations showed significant positive enrichment in the remission group (Figure 1c). Together, these data show that epithelial gene expression is associated with clinical remission to etrolizumab.

### Systems genomics links patient genotype to perturbed epithelial gene regulatory networks

Given the importance of gene sets within the epithelial compartment for achieving clinical remission to etrolizumab, we next characterised potential upstream genetic drivers contributing to perturbed epithelial gene regulatory networks in UC patients. To evaluate this, we first performed TF activity inference from fold change in the epithelial compartment of UC patients vs non-IBD controls in the scIBD resource [19], and then applied iSNP [14] to model how non-coding SNPs in etrolizumab-treated UC patients from the HICKORY cohort may perturb these TFs (Figure 2).

**Figure 2.**
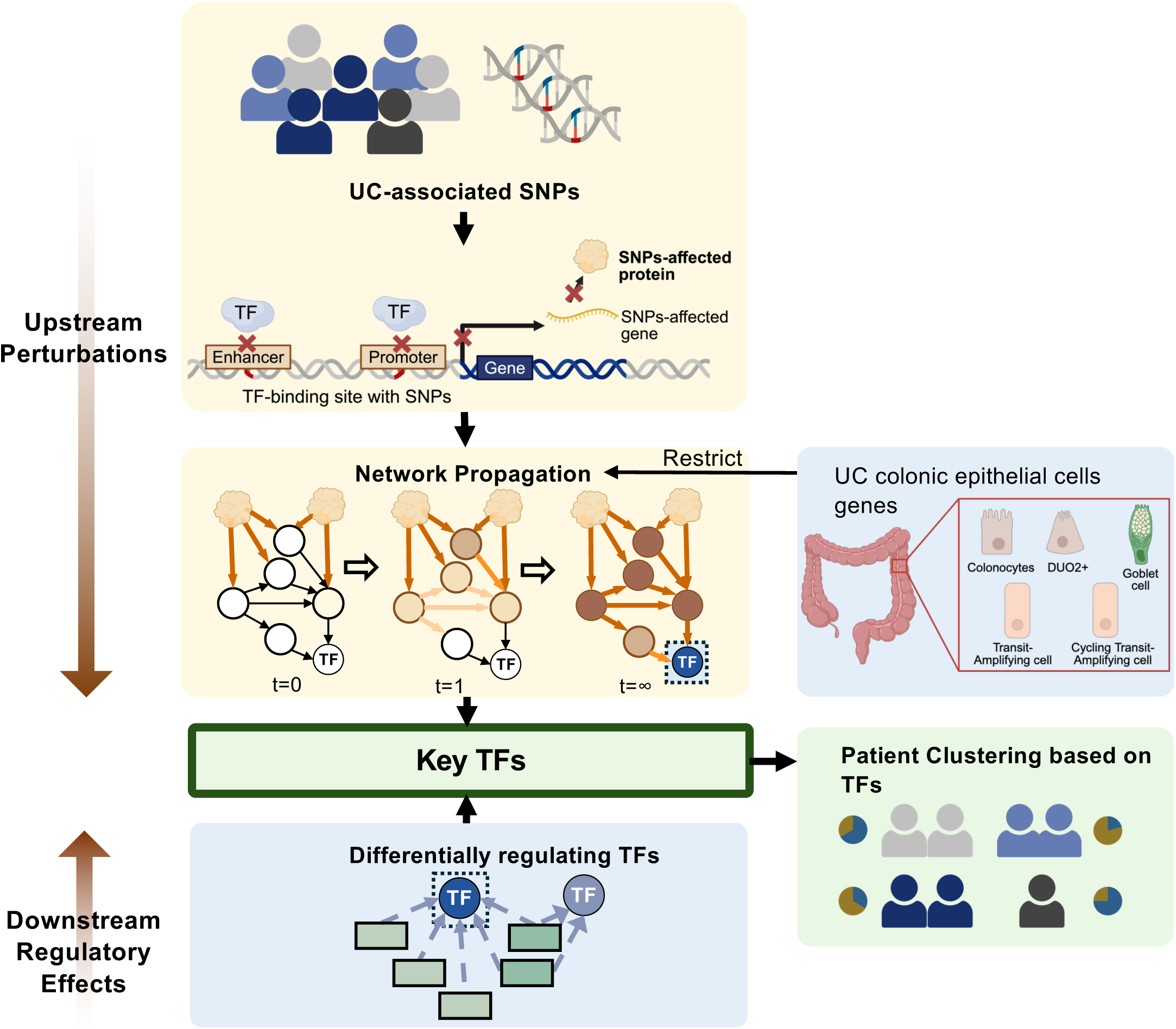
**Overview of the Workflow**. Ulcerative colitis (UC)-associated single nucleotide polymorphisms (SNPs) were extracted from UC patients in the HICKORY cohort. Based on genomic annotations indicating whether the SNPs were located in enhancer or promoter regions, we identified the corresponding SNP-affected genes and their protein products; Network heat propagation was then applied to assess the impact of non-coding SNPs within the signalling network, restricted to genes expressed in colonic epithelial cells from UC patients, and SNP-propagated transcription factors (TFs) were identified from the resulting network; TFs with differential regulatory activity were inferred from the scIBD single-cell expression data, and those overlapping with the SNP-propagated TFs were retained; Patient-specific profiles of overlapping TFs were used to cluster individuals, and clinical outcomes were compared between clusters.

Using epithelial cell scRNAseq data from the scIBD resource, TF activity in epithelial cells was inferred using decoupleR [21] from DEGs, identifying 554 TFs showing differential regulatory activity in total across all epithelial cell types, with the following number of differentially active TFs in each epithelial cell subtype: 196 (all epithelial cells), 97 (TA cells), 15 (adult colonocytes), 27 (goblet cells), 198 (cycling TA cells) and 21 (DUOX2+ epithelial cells).

Using iSNP, we then examined how UC-associated regulatory genetic variants may influence signalling in colonic epithelial cells and ultimately converge on differentially active TFs in each epithelial cell type. From the HICKORY cohort, 34 UC-associated non-coding SNPs were mapped to 87 proteins that may be affected by these variants. Using scRNAseq data from the scIBD resource, we identified genes that were consistently expressed across epithelial cell types and used these to refine the signalling network to include only interactions that occur in epithelial cells. By applying network propagation on this epithelial-specific signalling network, we were able to trace how regulatory genetic changes may spread through epithelial signalling pathways starting from UC SNP-affected proteins. This analysis identified the TFs most likely to be perturbed by UC-associated non-coding SNPs for each epithelial cell type, providing insights into cell type-specific regulatory effects (Figure 3). In all epithelial cells, 18 TFs were predicted to be perturbed, with affected pathways including GPCR, FOXO, Notch and RHO signalling, as well as cell cycle regulation. At the individual cell type level, colonocytes showed perturbation of 5 TFs, with disruption of RHO signalling. In transit amplifying (TA) cells, 13 TFs were affected, with mTOR and RHO signalling identified as the main affected pathways. DUOX2+ epithelial cells showed a broader pattern of signalling disruption, with 14 TFs affected through toll-like receptor, RHO and cytokine signalling pathways. In goblet cells, 12 TFs were predicted to be perturbed, primarily via disruption of Notch signalling. Cycling TA cells exhibited the greatest number of affected TFs, with 21 TFs perturbed across multiple pathways, including RHO and Notch signalling, stress response, and FOXO-regulated transcription.

**Figure 3.**
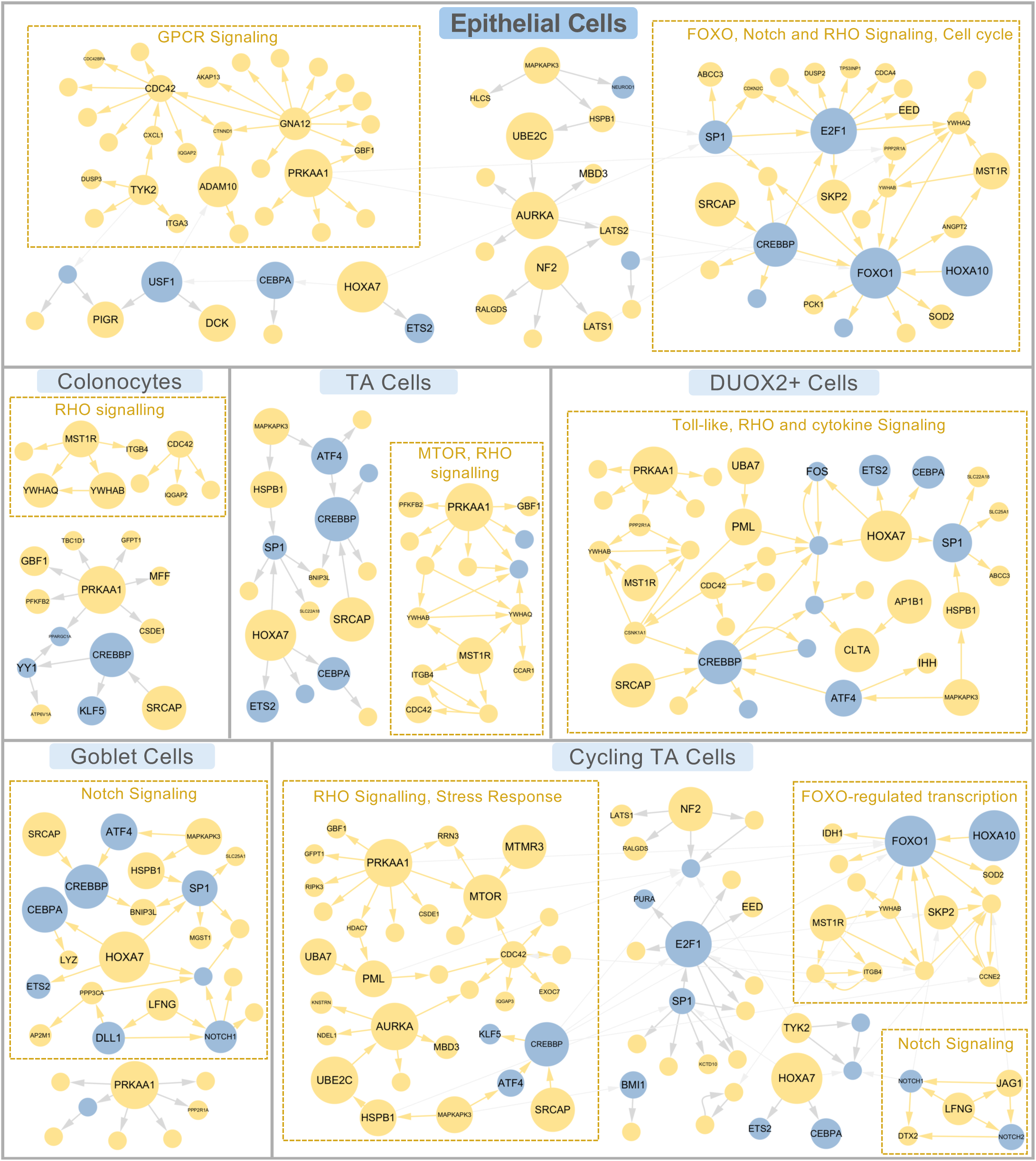
Ulcerative colitis-associated non-coding SNPs affect RHO signalling, Notch signalling, FOXO-regulated transcription, and other pathways in epithelial cells. SNP-propagated signalling network in all epithelial cells, goblet cells, DUOX2+ cells, colonocytes, cycling TA cells and TA cells, grouped by Markov Clustering (MCL). The modules are functionally annotated using Reactome enrichment, with each yellow box denoting a MCL module and its associated enriched pathways. Node size reflects the number of patients in whom that protein is affected by SNPs. Protein names are displayed for proteins affected in more than five patients. Networks with fewer than three nodes are excluded from the visualisation.

There were 23 TFs that overlapped between TFs inferred from differential expression and SNP-perturbed TFs. 14 TFs remained after filtering for TFs that were affected in more than 10 etrolizumab-treated UC patients in the HICKORY cohort (Figure 4).

**Figure 4.**
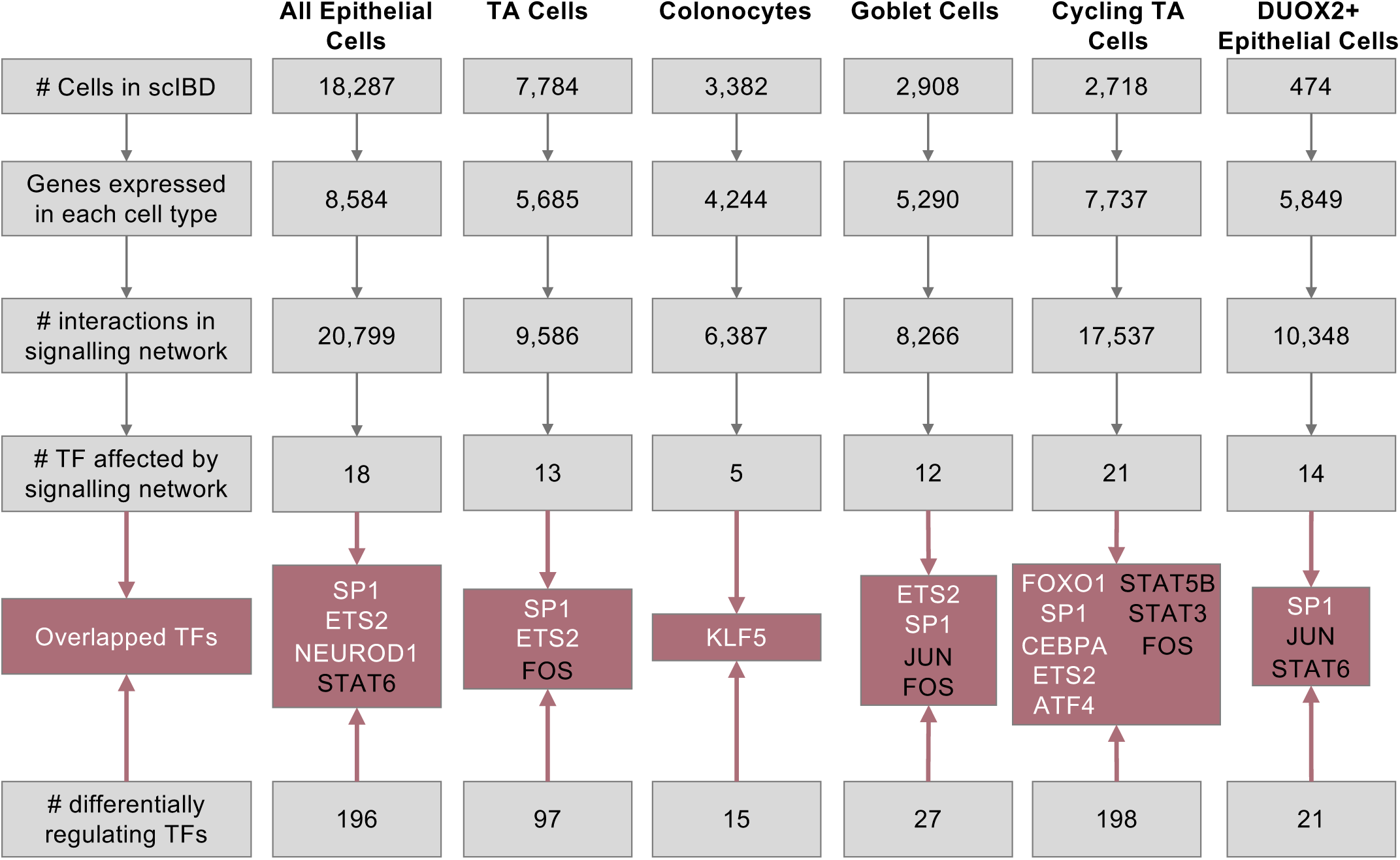
Cell type-specific network filtering and the resulting overlapping transcription factors. The number of cells and genes expressed in each selected cell type in scIBD dataset; the number of edges (protein-protein interactions) in OmniPath after filtering; the number of TFs affected after network propagation; the number of predicted differentially active TFs; and the overlapping TFs between the SNP-perturbed TFs and the differentially active TFs are shown. TFs indicated in white denote those retained for subsequent clustering analysis, whereas TFs indicated in black were excluded because they were affected in less than 10 etrolizumab-treated UC patients.

### SNP-propagated epithelial gene regulatory networks stratify UC patients into distinct clusters with differential treatment response to etrolizumab

We constructed patient-specific profiles based on these 14 TFs that were predicted to exhibit altered activity in epithelial cells and to be perturbed by non-coding IBD-associated SNPs. Clustering analysis of patient-specific gene regulatory networks revealed 6 patient clusters (Figure 5). Using a threshold of 25% of patients affected within a cluster, each cluster was characterised by a distinct combination of perturbed TFs across epithelial cell types (Figure 6a). Cluster 1 was characterised by perturbation of ETS2 in TA cells, KLF5 in colonocytes, and SP1 in DUOX2+ epithelial cells. Cluster 2 was distinguished by affected NEUROD1 in epithelial cells, ATF4 and CEBPA in cycling TA cells, and SP1 in both goblet and DUOX2+ epithelial cells. Cluster 3 showed the broadest pattern of SP1 perturbation, spanning epithelial, goblet, cycling TA, and DUOX2+ epithelial cells, alongside ETS2 disruption in epithelial, TA, and cycling TA cells, and CEBPA perturbation in cycling TA cells. Cluster 4 was characterised by SP1 perturbation in epithelial, goblet, and DUOX2+ epithelial cells, ETS2 in TA cells, CEBPA in cycling TA cells, and KLF5 in colonocytes. Cluster 5 was distinguished by ETS2 perturbation in epithelial cells, SP1 in TA, goblet, and cycling TA cells, ATF4 in cycling TA cells, and SP1 in DUOX2+ epithelial cells. Cluster 6 was characterised by ETS2 perturbation in epithelial and goblet cells, alongside SP1 and ATF4 disruption in cycling TA cells. Notably, FOXO1 in cycling TA cells was affected in nearly all patients across clusters, with 430 out of 452 patients affected, suggesting a shared regulatory disruption independent of cluster assignment.

**Figure 5.**
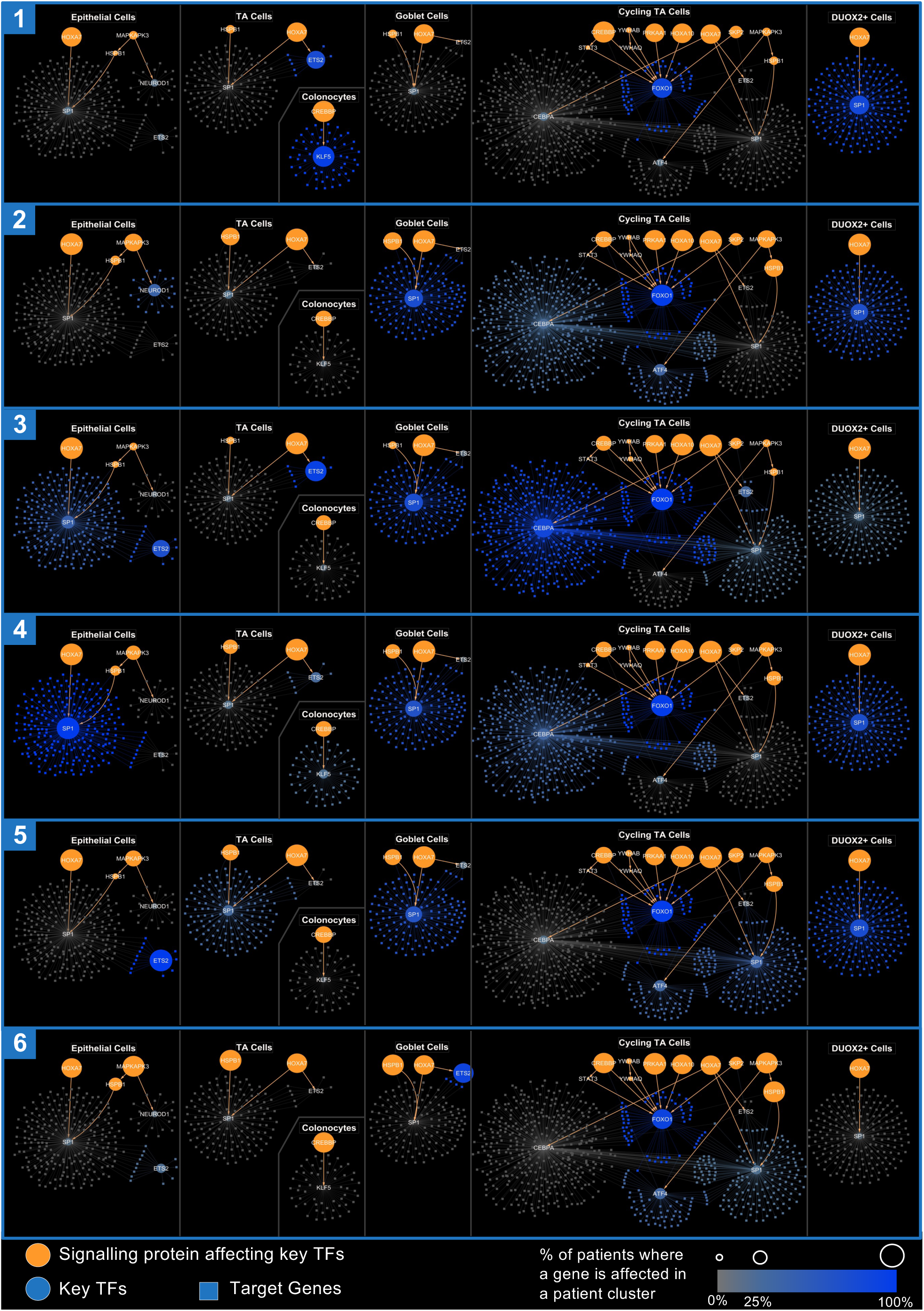
Cluster-representative gene regulatory networks across the colonic epithelial compartment and its subtypes. Networks are shown for the overall epithelial compartment and for five epithelial cell subtypes: transit-amplifying (TA) cells, adult colonocytes, goblet cells, cycling TA cells, and DUOX2+ epithelial cells. Orange circles represent signalling proteins upstream of the key transcription factors (TFs), blue circles represent the key TFs themselves, that is the overlapping TFs retained for clustering, and blue squares represent target genes of those TFs. Node size reflects the proportion of patients in the cluster in whom the corresponding protein was SNP-propagated. TFs and target genes are shown in blue where this proportion exceeds 25%, with darker shading indicating a higher proportion, and in grey where it is 25% or below. Edges represent signalling interactions from OmniPath and TF to target-gene relationships from the DoRothEA.

**Figure 6.**
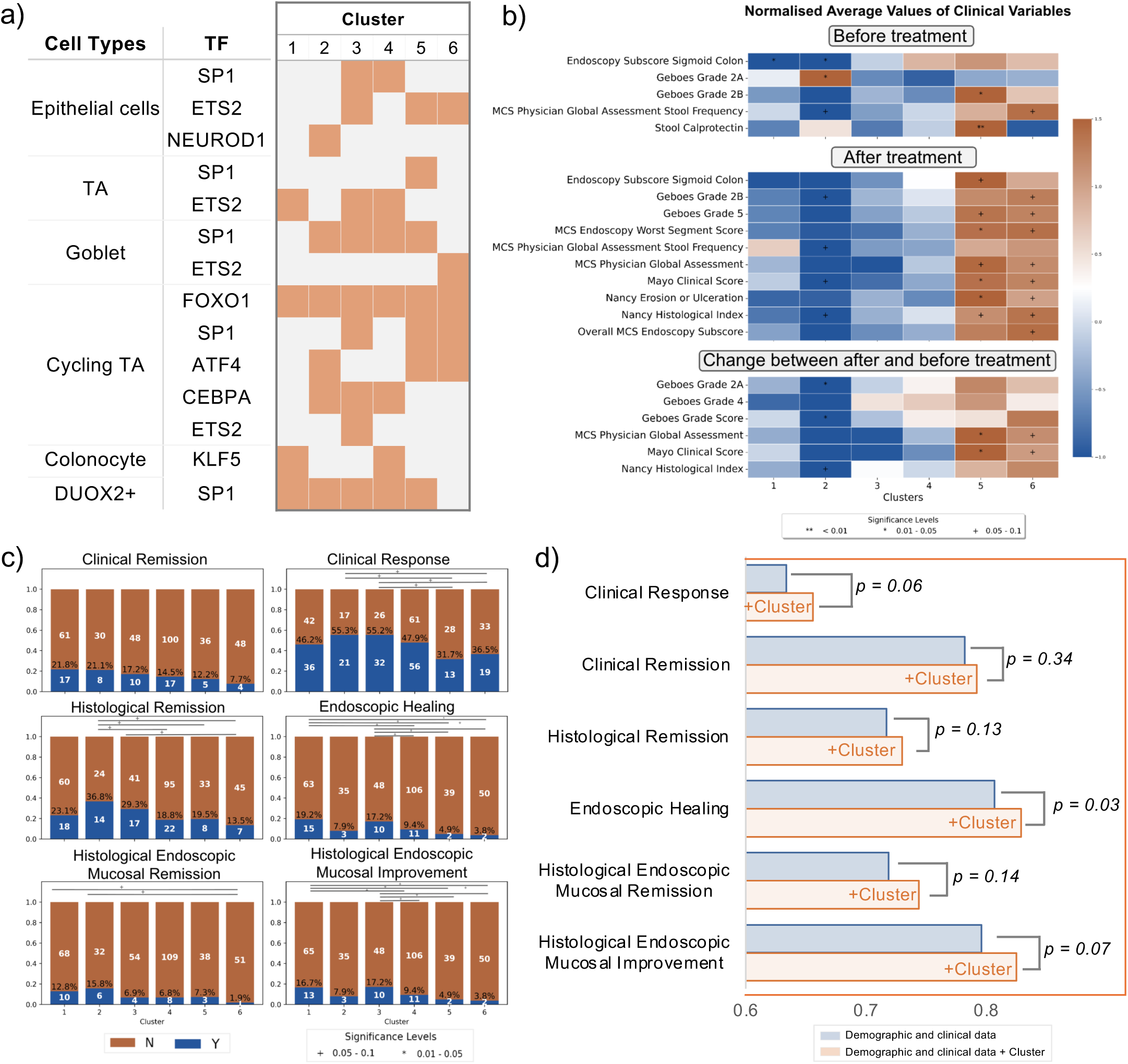
Patient clusters defined by SNP-propagated transcription factors associate with clinical features and etrolizumab response in UC. a) Cluster-representative SNP-propagated TFs across cell types. Orange indicates that more than 25% of patients in a cluster with a SNP-propagated TF, while light grey indicates a proportion of 25% or below. b) Normalised average values of clinical variables for etrolizumab-treated patients. Variables are grouped into three categories: measurements taken before treatment (top), after treatment (middle), and the change between the two time points (bottom). Statistical comparisons across clusters were performed as follows. Binary features: Chi-squared test; ordinal variables: Mann–Whitney U test; continuous variables: one-way ANOVA (normal distribution per Shapiro–Wilk) or Kruskal–Wallis test. All p-values were BH-corrected (threshold: 0.1). Statistical significance was determined using adjusted p-values. c) The proportion of patients achieving clinical remission, clinical response, histological remission, endoscopic healing, histological and endoscopic mucosal remission, and mucosal improvement across clusters. The proportional z-test was used as the statistical method, corrected using BH. d) Area under the receiver operating characteristic curve (AUROC) for predicting clinical endpoints. 2 Models were fitted and compared using a likelihood ratio test, with p-values derived from a chi-squared distribution based on the difference in degrees of freedom. Blue bars represent predictions based on demographic and clinical data alone, while orange bars represent predictions incorporating cluster information.

By analysing the clinical variables of patients who received etrolizumab, we next assessed whether patient clusters derived from SNP-propagated gene regulatory networks reflected heterogeneity in disease severity and clinical outcomes following etrolizumab treatment. 36 clinical variables were analysed at baseline, after the induction period (week 14), and as changes from baseline. Pairwise comparisons between the six patient clusters revealed distinct patterns. Patient cluster 1 showed the lowest disease severity scores (i.e. histological/endoscopic/clinical disease severity) while patient clusters 5 and 6 had higher disease severity scores at both baseline and week 14 and exhibited positive normalised changes from baseline, in contrast to the negative changes observed in cluster 2 (Figure 6b). No statistically significant differences between clusters were observed in clinical variables measured after the maintenance period (week 66), which may be attributable to the limited number of patients who completed the maintenance period (n = 63) and the limited availability of these clinical data.

Comparing the proportion of patients achieving several clinical endpoints revealed that patient clusters 5 and 6 exhibited the lowest proportions of remitters and responders, with reduced rates of histological remission, endoscopic healing, mucosal healing, HEMI, and HEMR (Figure 6c). In contrast, clusters 1 and 2 had higher proportions of patients achieving clinical remission, while clusters 1 and 3 showed higher clinical response rates. The proportion of patients achieving endoscopic healing was significantly higher in cluster 1 compared to cluster 6 (*p* = 0.04). Other inter-cluster differences were on the threshold of significance (*p* = 0.05–0.10), likely due to low power of the individual clusters. Comparing the proportion of patients achieving clinical response or remission between those who received etrolizumab and those who received placebo did not yield a statistically significant difference, which is again likely attributable to the limited sample size within individual clusters (Supplementary Figure 1).

To assess the added predictive value of this systems genomics-based patient clustering approach, we compared logistic regression models with and without cluster information across several clinical endpoints (Figure 6d). Adding genotype-driven cluster information to demographic and clinical data at baseline significantly improved the prediction of endoscopic healing (*p*=0.03). For clinical response and histological endoscopic mucosal improvement, the models with genotype-driven cluster information showed a trend toward better performance (*p*=0.06 and *p*=0.07, respectively). Adding cluster information did not meaningfully improve prediction of histological endoscopic mucosal remission or clinical remission (*p*=0.14 and *p*=0.34, respectively).

## Discussion

The marked pathogenic heterogeneity of ulcerative colitis (UC) is a major factor underpinning variable responses to treatments amongst the patient population [22]. Despite initial hopes that patient genotype could be harnessed to dissect this heterogeneity, mechanistically informative genotype-based patient stratification has not yet been realised in IBD or other complex diseases. In this study, we address this gap by analysing genotype data from the HICKORY phase III clinical trial of etrolizumab in UC and applying an established systems-genomics framework [14] that we contextualised here to the intestinal epithelial compartment.

Given that etrolizumab exerts key effects at the intestinal epithelial compartment [15], the known enrichment of UC-associated SNPs in the epithelial layer [16], and the importance of impaired epithelial barrier function as a central feature of UC pathogenesis [17], we posited that integrating patient genotype within an epithelial context would yield biologically meaningful stratification and provide novel insights into differential treatment outcomes. To test this, we performed a patient-specific integrative analysis of non-coding disease-associated GWAS variants with epithelial-specific signalling and gene-regulatory networks. This analysis identified 14 TFs predicted to be perturbed by SNP-propagated networks in epithelial cells in the HICKORY RCT cohort. Clustering 452 trial participants based on these TF perturbations revealed six patient clusters that differed in baseline disease severity and etrolizumab treatment outcomes, suggesting epithelial regulatory programmes act as an important substrate for UC heterogeneity and treatment variability.

Several of the TFs identified in our analysis have biological roles consistent with IBD pathophysiology. ETS2 has recently been identified as a key driver of intestinal inflammation in the context of Crohn’s disease, acting via inflammatory macrophages [23]. Our findings suggest that it may also have a key role in UC and may exert effects within epithelial cells. SP1, another key TF identified in our analysis, regulates the transcription of various genes involved in inflammation and epithelial processes in the intestine [24]. NEUROD1, though not previously associated with UC, plays an important role in intestinal epithelial cell differentiation by promoting enteroendocrine cell differentiation, whilst suppressing Paneth cell [25] and goblet cell lineages [26]. KLF5, another TF identified in our analysis, is critical for promoting epithelial cell proliferation [27] and has been shown to accelerate colitis development following specific deletion in murine models [28]. CEBPA is upregulated in UC patients [29], but its role in pathogenesis remains unclear. In colorectal cancer, CEBPA regulates epithelial proliferation, immune-related signalling, and epithelial–mesenchymal transition, suggesting that it may influence epithelial cell behaviour and inflammation in UC. ATF4 levels are significantly decreased in inflamed intestinal mucosa from patients with active UC [30]. ATF4 regulates genes involved in the inflammatory response [31], endoplasmic reticulum stress [32], autophagy [33], and amino acid metabolism [34], processes that are relevant to UC development. In murine models, ATF4 deficiency promotes intestinal inflammation by reducing glutamine uptake and antimicrobial peptide expression [30]. FOXO1 is upregulated in UC intestinal tissue [35] and is involved in apoptosis, stress responses, DNA damage repair, and metabolism [36]. In mucin-deficient colitis models, FOXO1 helps maintain epithelial barrier function [37]. Downregulation of FOXO1 reduces activation of the TLR4/MyD88/MD2–NF-κB inflammatory pathway, decreases mucosal barrier permeability, and upregulates tight junction proteins [35]. Collectively, the biological functions of these TFs support the concept that genetically perturbed epithelial regulatory networks could influence both disease severity and different outcomes to etrolizumab treatment.

Previous genetic studies of UC have been limited in their capacity to move beyond mapping non-coding risk to loci [38], bulk eQTLs [39], or protein-protein interaction networks [13]. Our approach links variants to epithelial signalling cascades and affected TFs at the patient level, then cross-validates with TF activity inferred from colonic epithelial gene expression. Importantly, clustering based on these epithelial-contextualised gene regulatory networks exhibited differences in UC disease severity and differential outcomes to etrolizumab therapy. Notably, patient cluster 1 showed the lowest baseline disease severity scores across histological, endoscopic, and clinical measures, whereas clusters 5 and 6 showed higher baseline severity. Furthermore, clusters 5 and 6 also demonstrated lower remission rates at week 14 of 7.7% and 12.2%, respectively, compared with over 20% in clusters 1 and 2. Incorporating genotype-derived cluster information into logistic regression models containing clinical data significantly improved prediction of endoscopic healing (p=0.03), with trends observed for predicting clinical response and histological endoscopic mucosal improvement (HEMI). Overall, these findings indicate that genotype data, when applied through the lens of systems genomics, can yield clinically and mechanistically meaningful patient stratification.

This approach has important translational implications for precision medicine in UC. It could result in the identification of patients who are more likely to respond to treatments from the outset, which could reduce the use of ineffective treatments with potential adverse effects in those unlikely to experience benefit. Furthermore, treatments that may appear ineffective in heterogeneous trial populations could demonstrate efficacy within molecularly defined patient subgroups, thereby increasing the likelihood of successful drug development. This is particularly relevant given that approximately 70% of UC patients still fail to achieve sustained clinical remission [2], and 14% ultimately require colectomy despite the availability of multiple advanced therapies [40]. Moreover, such an approach could enable the development of precision clinical trials. Focusing on genotype is particularly practical in this context, given that it is stable over time, relatively inexpensive to measure compared with metabolomics and transcriptomics, and readily scalable across large cohorts to inform effective patient stratification.

This study had a number of strengths. Clustering patients based on genotype and linking this to a rigorously phenotyped phase III clinical trial cohort enabled us to relate underlying genetic architecture to treatment response. The integration of genomic and transcriptomic information provided a multi-layered view of biological mechanisms that drive these differences. In addition, cross-validation of TFs using both iSNP and TF activity inference increased confidence in the identified regulatory programmes. Furthermore, the use of publicly available UC single-cell transcriptomics datasets enabled cell type–specific analyses despite the absence of patient-derived single-cell data in the HICKORY trial, increasing the biological resolution of our findings.

There were several limitations to the current investigation. The genotype-driven epithelial gene regulatory networks identified in this analysis were based on computational modelling of genotype and transcriptomics data, which require further experimental validation, but this was considered outside the scope of the study. Furthermore, the network propagation relied on a curated interaction network, which may be subject to literature bias towards well-studied genes and pathways. Additionally, cluster definitions are influenced by the chosen thresholds and distance metrics, which introduce some inherent subjectivity. Despite improving prediction of certain outcome measures, genotype-based clustering did not significantly improve prediction of clinical remission or histological endoscopic mucosal remission, which may reflect the low effect size of the drug, limited statistical power of the individual patient clusters, or other contributing factors beyond genetic architecture. The ability to assess long-term (week 66) treatment outcomes was also constrained by the limited number of patients completing the maintenance period (n=63). Finally, the current study evaluated responses only to etrolizumab and has not yet been evaluated for other IBD therapies.

Importantly, the analytical framework presented in this study is flexible and can be readily adapted to other complex diseases influenced by non-coding genetic variation. By linking patient-specific genetic variants to perturbed signalling and gene regulatory networks in disease-relevant cell types, this strategy provides a generalisable approach for translating genetic variation to functional biological insights. Such frameworks could provide novel insights for other complex diseases where specific cellular compartments are central to pathogenesis, including rheumatoid arthritis, type 2 diabetes, and asthma. Furthermore, integrating systems genomics modelling into clinical trial design could facilitate precision clinical trials and accelerate drug development by identifying molecularly defined patient subgroups most likely to benefit from specific therapies.

In summary, by harnessing a multi-layered systems genomics framework, we reveal epithelial-specific gene regulatory networks that capture UC heterogeneity in a large phase III clinical trial patient cohort, providing novel insights into genotype-driven epithelial mechanisms associated with differential treatment outcomes to etrolizumab. Collectively, this work highlights the potential for harnessing genotype for biologically informed patient stratification, which could be leveraged for future precision medicine strategies in IBD.

## Methods

### Differential expression analysis and cell deconvolution analysis

Colonic tissue bulk transcriptomic profiles from etrolizumab-treated UC patients in the phase III HICKORY trial cohort (n = 514) at baseline (week 0) and post-induction (week 14) were analysed. For each time point, differential expression between clinical remission and non-remission was assessed with DESeq2 [41]. P-values were adjusted for multiple testing using the Benjamini–Hochberg (BH) method, and genes with absolute log2 fold-change > 1 and adjusted p-value < 0.01 were defined as differentially expressed.

To relate transcriptional differences to specific colonic cell populations, we mapped differentially expressed genes to reference cell-type signatures derived from the single-cell IBD (scIBD) dataset [19]. Enrichment of differentially expressed genes (DEGs) in each scIBD major cell type (epithelial, mesenchymal, endothelial, B plasma, neural, CD4T, CD8T, ILC and myeloid cells) was tested using gene set enrichment analysis (GSEA) [42].

Bulk transcriptomic profiles from the HICKORY cohort were also deconvoluted using xCell [20]. Only cell types normally present in the intestine were retained (Supplementary Table 1). For each sample, we obtained xCell enrichment scores and then grouped samples by week 14 outcome (remission vs non-remission). StromaScores and ImmuneScores were calculated by averaging scores of stromal (e.g. endothelial cells, fibroblasts) and immune cell types, respectively. Differences in individual cell type scores and composite scores between remission and non-remission were assessed using t tests, and the resulting p-values were adjusted using the BH method to account for multiple testing. An adjusted *p* < 0.05 was considered statistically significant.

### Identifying UC SNP-affected proteins

Genotype data from 452 etrolizumab-and placebo-treated UC patients in the phase III HICKORY trial cohort were analysed using the Integrative SNP Network Platform (iSNP), which is described by Brooks-Warburton et al. [13]. Briefly, UC-associated SNPs were first identified from key GWAS studies [43,44] and annotated to determine which occurred within the enhancer or promoter regions of the genome. Promoter regions were defined as sequences spanning 5 kilobases upstream of the transcription start site or to the end of the first exon of the corresponding gene, based on the UCSC genome browser [45]. Enhancer regions were defined using the HEDD database (date of download: 13/08/2023) [46]. SNP selection was further filtered by retaining only those overlapping with Chromatin Immunoprecipitation Sequencing (ChIP-Seq) peaks identified within colonic tissues.

The functional impact of these non-coding SNPs on transcription factor binding sites (TFBS) was assessed by extracting ±50 bp flanking sequences and comparing ancestral and mutant alleles using two complementary in silico tools, RSAT matrix-scan and FIMO. Each non-coding SNP was classified as causing a gain, loss, or no change in TFBS, with only those predicted to cause a clear gain or loss retained for further analysis. Genes whose promoter or enhancer regions contained such variants were designated as ‘SNP-affected genes’, and the corresponding encoded proteins were considered ‘SNP-affected proteins’.

### Network propagation from UC SNP-affected proteins in colonic epithelial cells

To model the cumulative downstream effects of non-coding SNPs across the human cellular signalling network, network propagation was performed using the methodology described by Módos et al. [14]. Here, SNP-affected proteins were used as seed nodes from which heat-based network propagation (i.e. the directed version of the HotNet2 algorithm [47]) was implemented to identify downstream nodes perturbed within the human cellular signalling network. The signalling network was generated from OmniPath (as of October 2021) [48], comprising a directed giant component network with 6,954 protein nodes and 51,792 edges. To contextualise the signalling network to colonic epithelial cells, the iSNP pipeline was adapted by restricting network propagation to proteins expressed within colonic epithelial cells in UC patients. To do this, single-cell RNA-seq data from UC patients were obtained from scIBD (http://scibd.cn/) [19], which is a single-cell meta-analysis of IBD patients that combines highly curated single-cell transcriptomics datasets in a uniform workflow. The integrated gene expression matrix in H5AD format from scIBD was downloaded on 15 September 2024. The dataset was filtered to retain only epithelial cells from non-inflamed colonic tissues in adult UC patients. Genes were included if they were expressed in more than 30% of cells within any epithelial cell type, comprising transit-amplifying (TA), adult colonocyte, goblet, cycling TA, BEST4+ epithelial, DUOX2+ epithelial, enteroendocrine, tuft, fetal cycling TA, pediatric colonocyte, M-like, fetal progenitor, and Paneth cells, and if the overall gene count across all cells exceeded a threshold defined as the mean minus two standard deviations. Based on these genes, the OmniPath protein–protein interaction network was filtered to include only signalling interactions in which both source and target proteins were expressed in colonic epithelial cells in UC patients. Network propagation was then performed using this epithelial-specific signalling network, starting from the SNP-affected proteins. This allowed us to construct an overall epithelial compartment-specific SNP-propagated signalling network.

In addition, epithelial cell subtype-specific SNP-propagated signalling networks were constructed for epithelial cell subtypes relevant to IBD, including adult colonocytes, goblet cells, TA cells, cycling TA cells, and DUOX2+ epithelial cells. For each epithelial cell subtype, the signalling network from OmniPath was filtered to retain only genes expressed in more than 10% of the specific cell type. Network propagation was then run as described earlier for each epithelial cell subtype-specific signalling network.

In all instances, network propagation was run to convergence from SNP-affected proteins to downstream proteins including TFs. Following propagation, each protein in the network was assigned a Z-score reflecting the extent of its perturbation. TFs with a Z-score greater than 2 were designated as SNP-propagated TFs.

### Infer TFs with significantly altered activity from gene expression data

From the overall epithelial compartment-and epithelial cell subtype-specific SNP-propagated signalling networks, we identified SNP-propagated TFs (Z-score > 2). These were then compared to differentially active TFs inferred from expression data to identify overlapping TFs. To infer differentially active TFs in epithelial cells, the decoupleR Python package (version 1.6.0) [21] was used with the Univariate Linear Model (ULM) method, which infers TF activity from expression data. Log-transformed fold changes in gene expression between comparison groups served as input. P-values were adjusted using the BH method, with a significance threshold set at 0.05. This analysis was conducted for three comparison groups: (1) inflamed UC vs. non-inflamed UC, (2) inflamed UC vs. non-IBD control, and (3) non-inflamed UC vs. non-IBD control from the scIBD resource. TF activity inference was performed using both DoRothEA [49] and CollectTri [50] as reference regulon databases (downloaded 15 September 2024). The resulting sets of TFs with significantly altered activity were merged. Following this, we identified SNP-propagated TFs that overlapped with differentially active TFs inferred from the gene expression data.

### Clustering patients based on overlapping TFs

For each UC patient in the HICKORY cohort, a binary profile was constructed based on whether each overlapping TF was affected. Patients were then clustered based on these profiles. Pairwise Jaccard distances were calculated using the *pdist* function from SciPy (v1.10.1). The k-medoids clustering algorithm was applied, as it is more suitable for categorical data than k-means. Clustering was performed using the *scikit-learn-extra* package (v0.3.0). The optimal number of clusters was determined using both the silhouette score and the elbow method, as implemented in *scikit-learn*.

### Clinical feature comparison across patient clusters

The HICKORY phase III RCT had comprehensive and robustly characterised clinical parameters for each patient. To determine whether patient clusters exhibited distinct clinical characteristics, variables relating to clinical symptoms, biochemical disease activity, histopathological disease activity, endoscopic disease activity, composite disease activity, and prior treatment history were compared across three time points: baseline (prior to etrolizumab), week 14 post-etrolizumab, and week 66 post-etrolizumab (Supplementary Table 2). The change of clinical features from baseline was also compared. Statistical comparisons were performed based on the nature of each clinical feature. Binary features were analysed using the Chi-squared test with contingency tables. Ordinal variables were tested using the Mann–Whitney U test. For continuous variables, normality was assessed using the Shapiro–Wilk test. Normally distributed data were analysed with one-way ANOVA; otherwise, the Kruskal–Wallis test was applied. Multiple hypothesis testing was corrected using the BH method, with an adjusted p-value threshold of 0.1.

Eight clinical endpoints were used to assess differences in treatment responses to etrolizumab between patient clusters (Supplementary Table 3): clinical remission, clinical response, histological remission, histological improvement, endoscopic healing, mucosal healing, histological–endoscopic mucosal improvement (HEMI), and histological–endoscopic mucosal remission (HEMR) as defined in the HICKORY trial [10]. Pairwise comparisons of proportions between clusters were performed using the two-proportional z-test (one-sided), and p-values were corrected for multiple testing using the BH method.

To evaluate whether the addition of cluster information improved prediction of the clinical endpoint, we first fitted a logistic regression model including demographic variables (age, sex) and clinical variables (anti-TNF failure history, previous refractoriness to corticosteroids or immunosuppressants, Mayo Clinic score at baseline, treatment received, stool calprotectin, C-reactive protein). We then fitted a second logistic regression model that additionally included cluster membership. Models were fitted in Python using *statsmodels* and compared with a likelihood ratio test. *P*-values were obtained from a chi-squared distribution based on the difference in degrees of freedom. Predicted probabilities from each model were used to calculate the area under the receiver operating characteristic curve (AUROC) to compare discriminative performance.

## Data availability

The source data underlying the figures are provided with this paper. Access to individual patient-level data may be requested by qualified researchers through the clinical study data request platform (https://vivli.org/).

## Acknowledgments

The authors are grateful for the feedback and advice to current and past members of the Korcsmaros and Powell Labs.

The authors are grateful to Dr. Jacqueline McBride (Genentech, South San Francisco) for supervising the etrolizumab trial omics data collection and for facilitating data access and integration and overseeing transcriptomic data analysis and interpretation.

## Author contributions

The study was conceived and the methodology designed by Y.L., J.P.T., D.M., A.P., and T.K. Software was developed by Y.L. and B.B., with contributions from D.M. The manuscript was drafted by Y.L. and J.P.T., and critically reviewed and edited by B.B., N.P., D.M., A.P., and T.K. Figures and visualisations were prepared by Y.L. and T.K. The project was supervised by A.P. and T.K.

