## Supplementary material for "Epithelial regulatory networks link non-coding genetic variation to clinical heterogeneity in ulcerative colitis"

### **Supplementary materials**

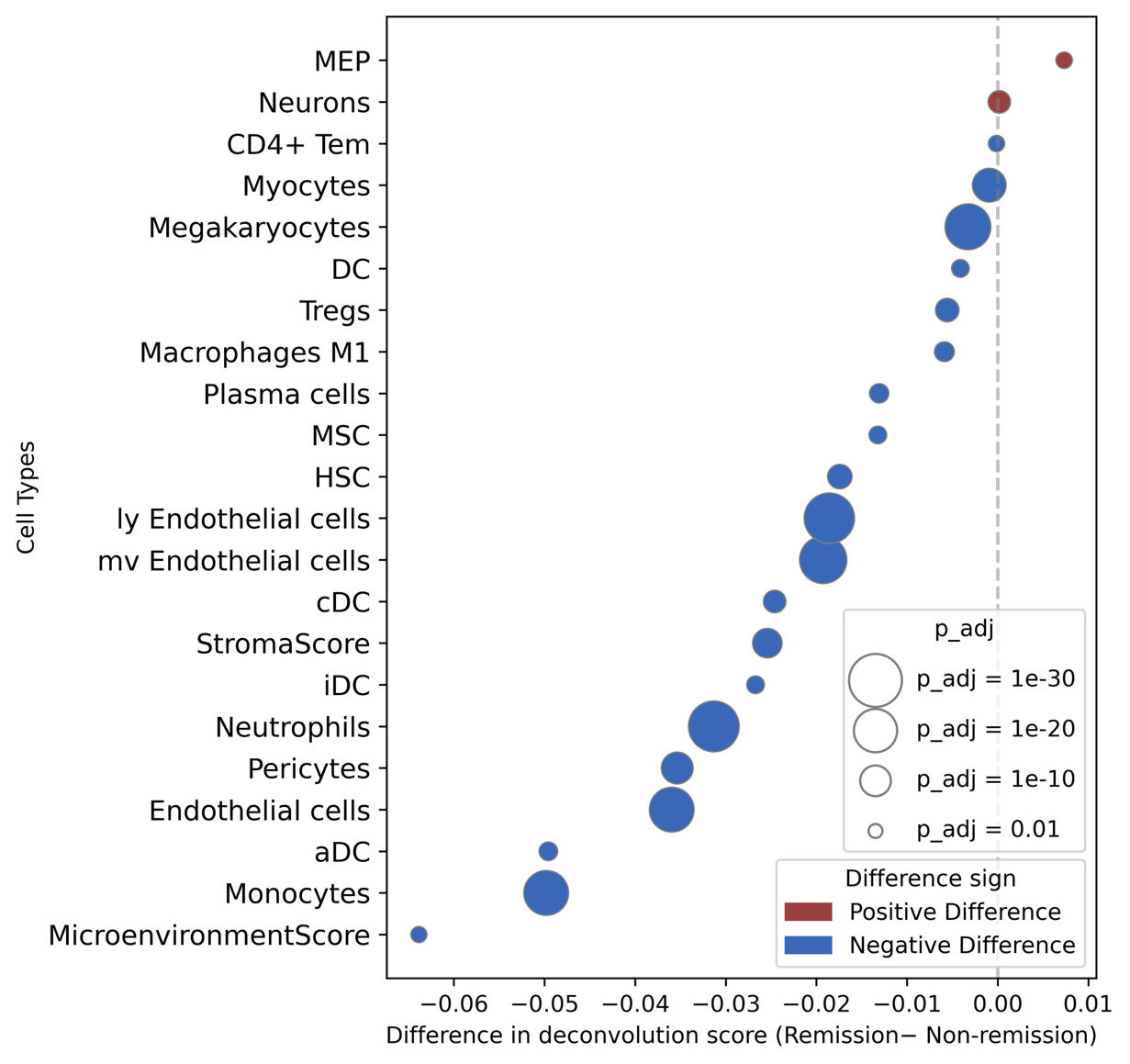

Supplementary Figure S1. Cell types with statistically significant differences in xCell cell deconvolution scores between remission and non-remission patients at baseline, restricted to *p*_adj_ < 0.05. The colour of each point denotes the direction of the difference (red for remssion and blue for non-remission s). Point size is proportional to -log_10_(*p*_adj_).

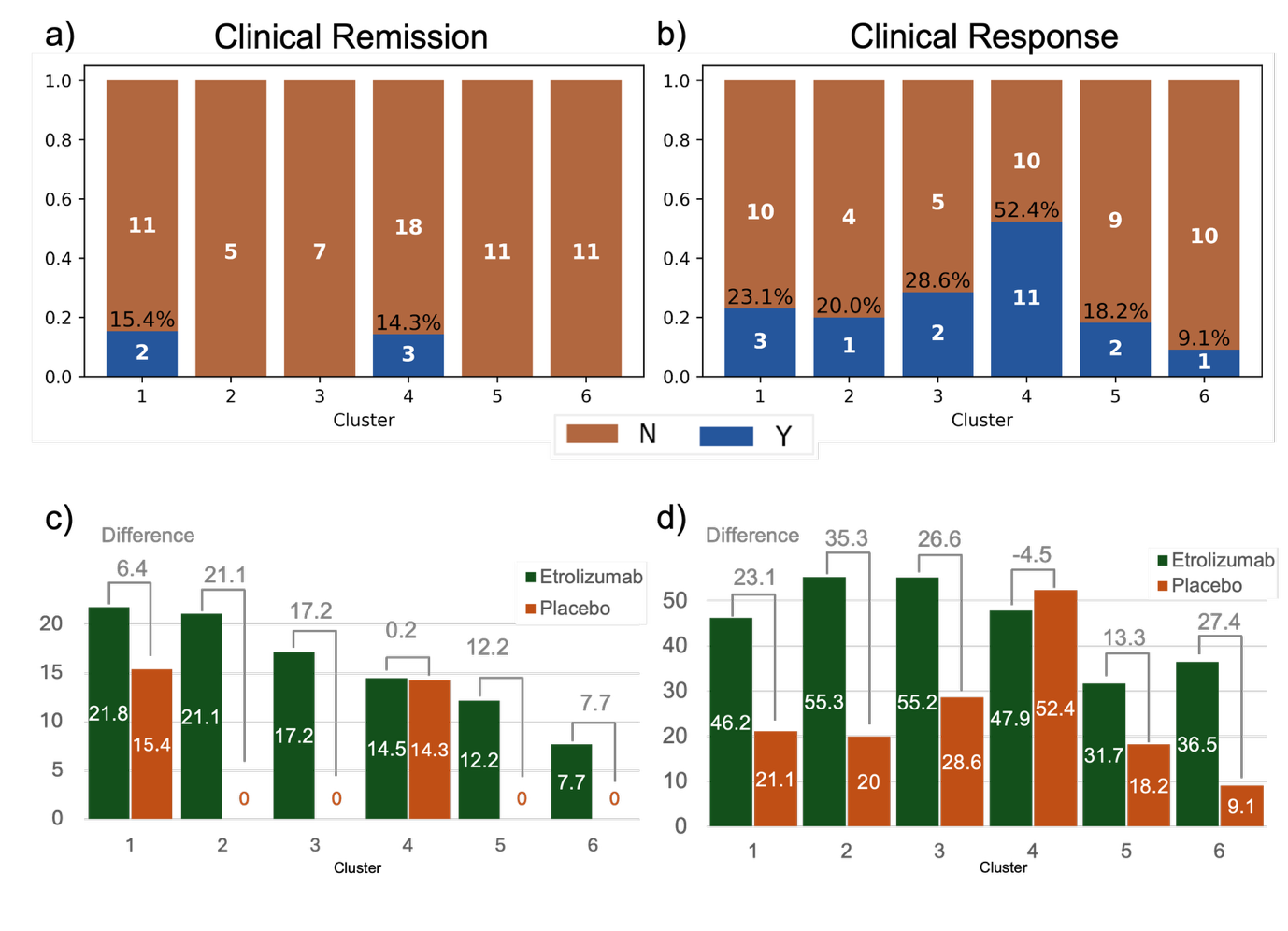

Supplementary Figure S2. The proportion of patients receiving placebo during the induction period achieving (a) clinical remission and (b) clinical response at week 14 across clusters. The proportion of patients achieving (c) clinical remission and (d) clinical response was compared between those receiving etrolizumab and those receiving placebo. The difference in proportion is annotated at the top.

| Cell Types in intestine | aDC, B-cells, Basophils, CD4+ memory T-cells, CD4+ naive T-cells, CD4+ T-cells, CD4+ Tcm, CD4+ Tem, CD8+ naive T-cells, CD8+ T-cells, CD8+ Tcm, CD8+ Tem, cDC, Class-switched memory B-cells, CLP, CMP, DC, Endothelial cells, Eosinophils, Epithelial cells, Erythrocytes, Fibroblasts, GMP, HSC, iDC, ly Endothelial cells, Macrophages, Macrophages M1, Macrophages M2, Mast cells, Megakaryocytes, Memory B-cells, MEP, Monocytes, MPP, MSC, mv Endothelial cells, Myocytes, naive B-cells, Neurons, Neutrophils, NK cells, NKT, pDC, Pericytes, Plasma cells, Platelets, pro B-cells, Smooth muscle, Tgd cells, Th1 cells, Th2 cells, Tregs |
| --- | --- |
| Cell Types that are not found in intestine | Astrocytes, Chondrocytes, Hepatocytes, Keratinocytes, Melanocytes, Skeletal muscle, Osteoblast, Adipocytes, Preadipocytes, Sebocytes, Mesangial cells |

Table S1. Cell types used in cell deconvolution analysis.

| Clinical Symptoms | Physician's Global Assessment  Rectal Bleeding  Stool Frequency |
| --- | --- |
| Biochemical Disease Activity | Albumin  C-reactive protein  Haemoglobin  Faecal Calprotectin |
| Endoscopic Disease Activity | Endoscopy Subscore Rectum  Endoscopy Subscore Sigmoid Colon  Endoscopy Subscore Desc Colon  Overall MCS Endoscopy Subscore  Endoscopy Worst Segment Score |
| Histopathological disease activity | Overall Geboes Grade Score  Geboes Grade 0: Structural (Architectural Change)  Geboes Grade 1: Chronic Inflammatory Infiltrate  Geboes Grade 2A: Eosinophils in Lamina Propria  Geboes Grade 2B: Neutrophils in Lamina Propria  Geboes Grade 3: Neutrophils in Epithelium  Geboes Grade 4: Crypt Destruction  Geboes Grade 5: Erosion or Ulceration  Nancy Histological Index  Nancy Acute Inflammatory Cells Infiltrate  Nancy Chronic Inflammatory Infiltrate  Nancy Erosion or Ulceration  Robarts Histopathology Index |
| Composite disease activity | Mayo Clinical Score |
| Prior Therapy | Prior corticosteroids dependency  Previously refractory to corticosteroids  Prior corticosteroids intolerance  Previously refractory to immunosuppressants  Prior immunosuppressants intolerance  Prior anti-TNF Non-Response  Prior anti-TNF Loss of Response  Prior anti-TNF Intolerance |

Table S2. Clinical features used to compare patient clusters.

| **Clinical Endpoints** | **Conditions** |
| --- | --- |
| Clinical Remission | Mayo Clinical Score at Week 14 ≤ 2, with individual subscore ≤ 1, rectal bleeding subscore = 0 |
| Clinical Response | Mayo Clinical Score with ≥ 3-point decrease  ≥ 30% reduction from baseline  ≥ 1 point decrease in rectal bleeding subscore or absolute rectal bleeding subcore = 0 |
| Histologic Remission | Geboes Grade Score at Week 14 < 2  Robarts Histopathology Index at Week 14 ≤ 3  Nancy Histological Index at Week 14 = 0 |
| Endoscopic Healing | Mayo Clinical Score at Week 14 ≤ 1 |
| Histologic endoscopic mucosal improvement | Mayo Clinical Score at Week 14 ≤ 1  Geboes Grade Score at Week 14  ≤ 3 |
| Histologic endoscopic mucosal remission | MCS Endoscopy Subscore at Week 14 = 0  Geboes Grade Score at Week 14  < 2 |

Table S3. 6 clinical endpoints and their corresponding conditions.
